# Auditory and Language Contributions to Neural Encoding of Speech Features in Noisy Environments

**DOI:** 10.1101/377838

**Authors:** Jiajie Zou, Jun Feng, Tianyong Xu, Peiqing Jin, Cheng Luo, Feiyan Chen, Jianfeng Zhang, Nai Ding

## Abstract

Recognizing speech in noisy environments is a challenging task that involves both auditory and language mechanisms. Previous studies have demonstrated noise-robust neural tracking of the speech envelope, i.e., fluctuations in sound intensity, in human auditory cortex, which provides a plausible neural basis for noise-robust speech recognition. The current study aims at teasing apart auditory and language contributions to noise-robust envelope tracking by comparing 2 groups of listeners, i.e., native listeners of the testing language and foreign listeners who do not understand the testing language. In the experiment, speech is mixed with spectrally matched stationary noise at 4 intensity levels and the neural responses are recorded using electroencephalography (EEG). When the noise intensity increases, an increase in neural response gain is observed for both groups of listeners, demonstrating auditory gain control mechanisms. Language comprehension creates no overall boost in the response gain or the envelope-tracking precision but instead modulates the spatial and temporal profiles of envelope-tracking activity. Based on the spatio-temporal dynamics of envelope-tracking activity, the 2 groups of listeners and the 4 levels of noise intensity can be jointly decoded by a linear classifier. All together, the results show that without feedback from language processing, auditory mechanisms such as gain control can lead to a noise-robust speech representation. High-level language processing, however, further modulates the spatial-temporal profiles of the neural representation of the speech envelope.

## Introduction

Speech perception is a complex process involving both auditory and language processing and language processing can feed back and modulate basic auditory perception (Ganong, 1980; Warren, 1970). A sound feature that strongly contributes to speech intelligibility is the speech envelope, i.e., slow fluctuations (< 16 Hz) in sound intensity (Drullman et al., 1994; Shannon et al., 1995). How auditory and language factors influence the neural processing of speech envelope has been extensively investigated but remains controversial. When listening to speech, neural activity tracking the speech envelope can be recorded either intracranially from auditory cortex (Nourski et al., 2009) or noninvasively by magnetoencephalography /electroencephalography (MEG/EEG) (Ding and Simon, 2012b). Even in noisy environments, neural tracking of the speech envelope remains robust as long as the speech stream is attended to (Ding and Simon, 2012a; Kerlin et al., 2010; Mesgarani and Chang, 2012; O’Sullivan et al., 2014; Zion Golumbic et al., 2013). Although it is well established that cortical activity can track the speech envelope, it remains controversial whether speech-tracking activity is generated by general auditory mechanisms or speech-specific neural computations (Ding and Simon, 2014).

One hypothesis is that envelope tracking responses are generated by non-speech-specific auditory mechanisms (Steinschneider et al., 2013), since the temporal envelope is a low-level acoustic feature well represented throughout the auditory system (Joris et al., 2004). Consistent with this hypothesis, envelope tracking responses can be seen in animals (David et al., 2009) and in humans listening to non-speech sound, e.g., amplitude modulated noise or tones (Lalor et al., 2009; Wang et al., 2012). Critically, some studies have found similar envelope tracking responses for intelligible speech and unintelligible speech such as time-reversed speech (Howard and Poeppel, 2010) and speech in an unknown language (Peña and Melloni, 2012). Based on the domain-general auditory encoding hypothesis, noise-robust neural tracking of the speech envelope can be explained by contrast gain control (Ding and Simon, 2013) or primitive auditory scene analysis (Bregman, 1990; Ding et al., 2014), i.e., sound source segregation based on acoustic features. Contrast gain control and primitive auditory scene analysis are general auditory mechanisms that have been observed in primary auditory cortex of animals (Micheyl et al., 2005; Rabinowitz et al., 2013; Rabinowitz et al., 2011).

Another hypothesis assumes that envelope tracking responses reflect interactions between auditory and language processing. Consistent with this hypothesis, some studies show that when speech is acoustically degraded to compromise intelligibility, neural tracking of the speech envelope shows reduced precision (Gross et al., 2013; Kong et al., 2015; Luo and Poeppel, 2007; Peelle et al., 2013). Furthermore, at the individual level, listeners showing more precise envelope tracking activity tend to understand speech better (Ding et al., 2014; Ding and Simon, 2013; Doelling et al., 2014). A potential concern about whether speech intelligibility directly modulates envelope tracking activity, however, is that intelligibility covaries with acoustic changes, task difficulty, top-down attention, and individual hearing functions, which are factors known to modulate envelope tracking activity (Kayser et al., 2015; Lakatos et al., 2013; Petersen et al., 2017).

Here, we investigate how auditory and language mechanisms separately contribute to envelope-tracking speech responses in noisy environments. Behaviorally, it is known that language information increases speech intelligibility in noise (Miller et al., 1951). Based on the domain-general auditory processing hypothesis, language knowledge facilitates speech recognition at a late stage, not reflected in the envelope tracking response. Based on the interactive processing hypothesis, however, language processing feeds back and modulate envelope-tracking activity. To distinguish these two hypotheses, we investigate the influence of language processing by comparing two groups of listeners, i.e., native listeners of the testing language and foreign listeners who do not understand the testing language. A low-level auditory task is employed to ensure attention, which does not require language comprehension. The speech signal is mixed with spectrally matched stationary noise at 4 signal-to-noise ratios (SNR), and we measure envelope-tracking activity from both groups of listeners using EEG.

## Materials and Methods

### Participants

Thirty-two right-handed adults participated in this experiment (18-29 years old; mean age, 22.9 years). All participants reported normal hearing and they were all undergraduate or graduate students from Zhejiang University. Sixteen participants (8 females) were native Cantonese listeners while the other 16 participants (8 females) were native Mandarin listeners who did not understand Cantonese. Participants were paid for their participation and the experimental protocol was approved by the Institutional Review Board of the Zhejiang University Interdisciplinary Center for Social Sciences. Informed consents were obtained from all participants.

### Stimuli and procedures

The speech recordings were selected from the fiction *Legends of the Condor Heroes*, narrated in Cantonese by a male speaker. One hundred and sixty sections were randomly selected from the fiction. Each section was 15 s in duration and was normalized to the same intensity, measured by the RMS. One hundred and twelve sections served as normal trials, while the other forty-eight sections served as outlier trials. In the outlier trials, two syllables were randomly selected and immediately repeated. Speech materials in the normal trials were checked to ensure there are no repeated syllables. For half of the outlier trials, the two syllables were repeated once and for the other half they were repeated twice. The syllable boundaries were manually determined.

Spectrally matched stationary noise was generated using a 12-order linear predictive coding (LPC) model estimated based on all the speech materials. The noise was mixed with speech at 4 signal-to-noise ratios (SNRs), i.e., +9 dB, −6 dB, −9 dB, and −12 dB. These 4 levels of SNRs were selected since a previous study has shown that the envelope-tracking neural responses were highly robust to noise when the SNR was above −6 dB. When the SNR dropped below −6 dB, however, the envelope-tracking response degraded. Here, +9 dB was chosen as a baseline and the other 3 SNR levels were selected to characterize how the envelope-tracking response was affected by the SNR.

Each SNR condition contained 40 trials, including 12 outlier trials. In the experiment, all 160 trials were presented in a randomized order. Background noise reduced the intensity contrast of speech, i.e., the standard deviation of the envelope divided by its mean, and distorted the spectrotemporal features, which was characterized by the correlation between the envelopes of the noisy stimulus and the underlying clean speech (Fig.1AB).

**Fig. 1.**
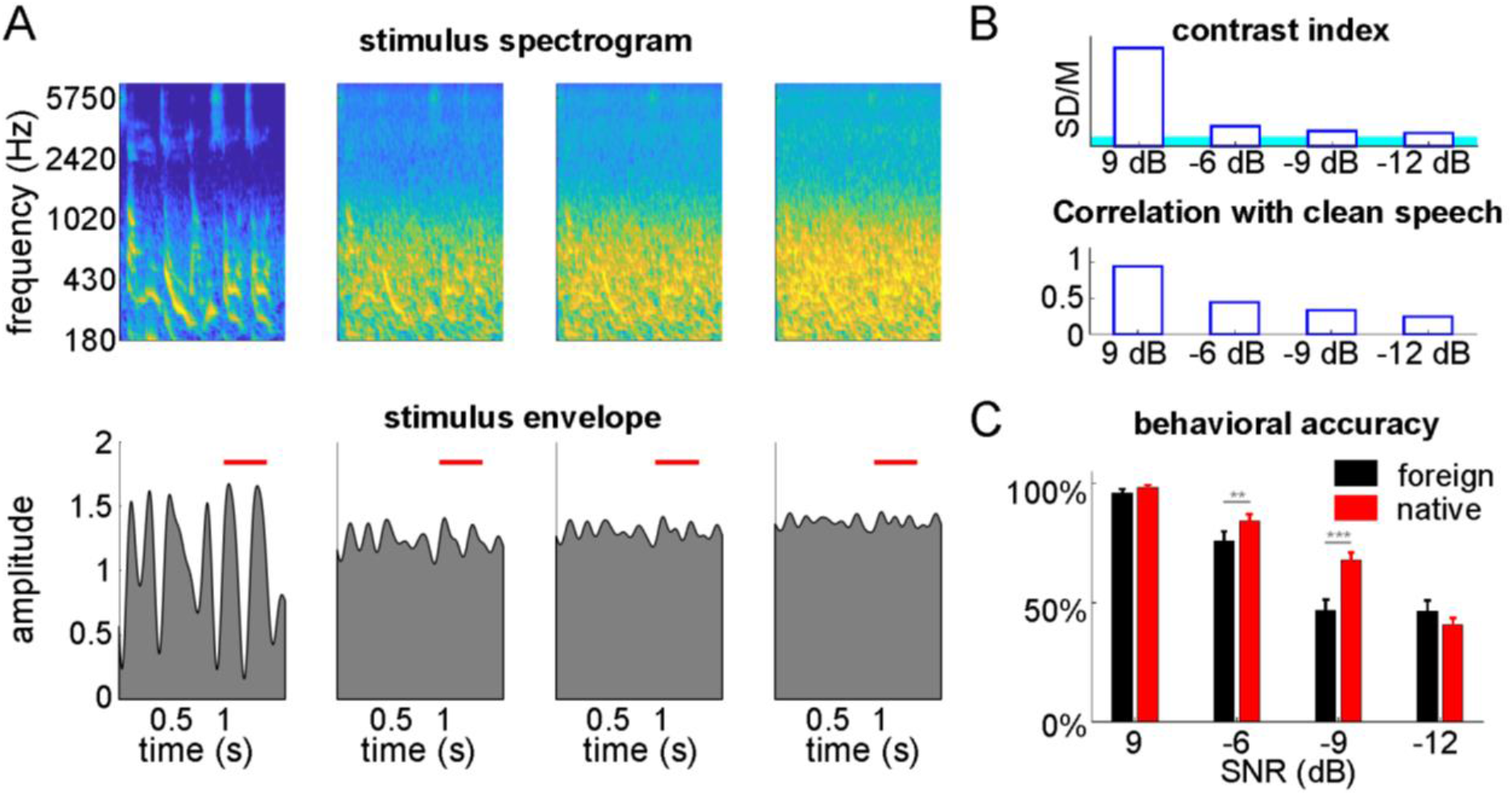
Stimuli and behavioral results. (A) Stimulus spectrogram (top) and envelope (bottom) under the 4 SNR conditions. The stimuli shown here consists of a repeated stimulus segment and the red lines illustrate the time interval when the segment repeats. (B) Contrast index of the stimulus (top) and the correlation between the stimulus envelope and the envelope of the underlying speech (bottom). The blue area covers the 95th percentile of the contrast index of stationary noise. (C) Behavioral results of native listeners (red) and foreign listeners (black) at each SNR. Error bars represent 1 SEM over listeners. **P<0.01, ***P<0.001 (bootstrap, FDR corrected).

After each stimulus, the listeners had to press 1 or 2 on the keyboard if they heard that a sound segment was repeated once or twice respectively. They had to press 0 if they did not hear any repeated segment. Since the number of normal trials was far greater than the number of outlier trials (28 vs. 12 trials), behavior accuracy would be high if listeners kept pressing 0. Therefore, in the following, behavior accuracy was characterized by analyzing the percent of outlier trials in which the participants made correct responses (i.e., true positive rate).

### EEG recording and preprocessing

EEG responses were recorded using a 64-channel Biosemi ActiveTwo system, sampled at 2048 Hz. Two reference channels were placed at the left and right mastoids respectively and four channels were used to record horizontal and vertical EOGs. The EEG signals were referenced offline by subtracting the average of the two mastoid recordings. EOGs artifacts were regressed out based on the least squares method (Ding et al., 2017). The EEG recordings and also the speech envelope were downsampled to 50 Hz and epoched based on the onset of each 15-s stimulus. The first 1 second of recording was removed to avoid the onset response. Only the normal trials (112 trials for each listeners) were analyzed, to avoid EEG responses evoked by the outliers, i.e., the repeated sound segments.

### EEG based envelope reconstruction

A linear decoder was used to reconstruct the temporal envelope of the underlying speech from the EEG response to a speech-noise mixture. The decoder applied a weighted average of the EEG response over time lags and channels using the following equation:

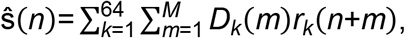

where *D_k_(n)* and *r_k_(n)* are the decoder weights and the EEG signal in channel *k* respectively. The order of the decoder, i.e., *M*, was 26, corresponding to a maximal time lag of 0.5 s. The decoder weights *D_k_(n)* were optimized so that the reconstructed envelope, *ŝ(n)*, approximated the underlying speech envelope. The decoder *D_k_(n)* was derived based on least-squares estimation with L2 regularization, i.e., normalized reverse correlation (Theunissen et al., 2001).

The neural reconstruction analysis was separately applied to each listener. The reconstruction accuracy was defined as the Pearson correlation between the reconstructed envelope and the envelope of the underlying speech. It was evaluated using 10-fold cross validation: Each time 90% of data was used to train the decoder and the rest 10% of data was used to evaluate the reconstruction accuracy. The procedure was repeated 10 times and the 10 reconstruction accuracy values were averaged. The regularization parameter for neural reconstruction was varied between 0 and 0.01 and the optimal value of 0.001 was chosen, which lead to the highest neural reconstruction accuracy averaged over conditions and participants.

To characterize the frequency bands in which the EEG response was most correlated with the speech envelope, envelope reconstruction was applied separately for bandpass filtered signals. In this analysis, a filter bank was used to decompose the audio and the EEG responses into narrow bands. The filter bank contained 25 filters with a bandwidth equal to 1 Hz, and the center frequencies linear increased from 1.5 Hz to 24.5 Hz in steps of 1 Hz. The frequency range in which the reconstruction accuracy is significantly higher than chance falls in the delta (1-4 Hz) and theta (4-8 Hz) bands (not shown), consistent with the literature (Ding and Simon, 2012b; Luo and Poeppel, 2007). Therefore subsequent analyses were restricted to 1-8 Hz.

A sigmoid function was used to characterize the relationship between behavioral accuracy and neural reconstruction accuracy, denoted as *y* and *x* respectively in the following. The sigmoid function, i.e., *y* = *A*+(1-*A*)/(1+exp(-*α*(*x*-*m*))), has 3 parameters, i.e., *A*, *α*, and *m*, which referred to the lower asymptote, the growth rate, and the location of this sigmoid function, respectively. The 3 parameters were fitted using the least squares method.

### Temporal response function

Neural reconstruction characterized how accurately the speech or stimulus envelope was represented in the brain by integrating EEG responses over time and channels. A neural encoding model, i.e., the temporal response functions (TRF), was used to further characterize the spatial and temporal patterns of the neural responses. The TRF could be formulated as the following:

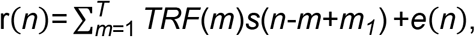

where *r(n)*, *TRF(n)*, *s(n)*, and *e(n)* was the EEG response, the TRF, the sound envelope, and the residual error respectively. The order of the TRF model, i.e., *T*, was 26, corresponding to 0.5 s. The integration process started from *m* = −5, so that the TRF contained a 5-sample, i.e., 0.1 s, prestimulus interval. The TRF was computed based on the least-squares estimation with L2 regularization. The regularization parameter was tuned to provide the highest predictive power, which was defined as the correlation between the predicted neural responses, i.e., 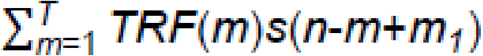, and the actual responses. Specifically, the regularization parameter was varied between 0 and 0.3 and the results showed that 0.18 was the optimal parameter that resulted in the highest predictive power averaged over all channels, conditions, and participants.

### Classification of the listeners and SNR

A classification analysis was employed to quantify whether the spatial and temporal profiles of the TRF were modulated by the language knowledge and the stimulus SNR. In this analysis, the 2 groups of listeners, i.e., native and foreign listeners, and the 4 levels of SNR were classified based on the TRF. The high-dimensional (time by channel) TRF was first reduced in dimension using the linear discriminant analysis (LDA), which projected high-dimensional data to dimensions in which data from different classes were maximally separated (Duda et al., 2012). A Euclidean distance-based classifier was then used to classify the listener groups and the SNR levels based on the first 2 LDA dimensions. The classifier calculated the center of each class based on the training set and each sample in the testing set was classified based on its distance to all the class centers. If a testing sample was close to the center of class A in terms of the Euclidean distance, it was attributed to class A. Since there were 16 listeners in each group, the classifier’s performance was evaluated using 8-fold cross validation: Each time, fourteen listeners were used to train a classifier while the other two listeners were used to evaluate classification accuracy.

When the LDA was applied to the time by channel 2-dimensional TRF (*T* x 64), the TRF was reshaped to a 1-dimensional vector (64*T* x 1) by concatenating the responses from different channels. The LDA feature vector was therefore also a 64*T* x 1 vector. To characterize the spatial and temporal profiles of the LDA feature vector, it is reshaped back to a time by channel 2-dimensional matrix and the spatial and temporal profiles were the first left and right singular vectors extracted by singular value decomposition (SVD).

### Statistical tests

The bootstrap significance test is based on a bias-corrected and accelerated procedure (Efron and Tibshirani, 1994). In this procedure, all participants were resampled 5000 times with replacement and each time the participants being sampled were averaged, resulting in 5000 mean values. For paired comparisons (e.g., comparisons between SNR conditions), bootstrap was applied to the difference between conditions, if *N_S_* out of the 5000 mean values were greater (or smaller) than 0, the significance level is *N_S_*/5000. For unpaired comparisons, i.e., comparisons between listener groups, data from the foreign listeners were resampled and used to estimate the null distribution. If the mean value over native listeners is greater (or smaller) than *N_S_* out of the 5000 resampled mean values of the foreign listeners, the significance level is *N_S_*/5000.

In the neural reconstruction analysis, chance-level reconstruction accuracy was estimated by constructing surrogate neural responses. A surrogate neural response was created by the circularly shifting the actual response by time lag T, which was between 100 s and 300 s in steps of 2 s. This procedure resulted in 101 surrogate neural responses. If the actual reconstruction accuracy was higher than the 95% percentile of the chance-level reconstruction accuracy, it was statistically significant (P < 0.05). A similar process was used to estimate the chance-level predictive power and RMS of the TRF. When multiple comparisons were involved, the p-value was further adjusted using the false discovery rate (FDR) correction.

## Results

### Behavioral results

In the experiments, listeners have to detect repeated sound segments in a continuous speech stream. The percent of repeated segments detected by native and foreign listeners are shown in Fig. 1C. Two-way repeated-measures ANOVA reveals a significant main effect of SNR (F_3,90_ = 180.81, P < 0.001) and a marginally significant main effect of listener groups (F_1,30_ = 3.74, P = 0.063). The interaction between the two factors also reaches significance (F_3,90_ = 9.91, P < 0.001). Additionally, the behavioral accuracy is significant higher for native listeners at −6 dB (P = 0.007, bootstrap, FDR corrected) and −9 dB (P < 0.001, bootstrap, FDR corrected). These behavioral results suggest that language knowledge facilitates the detection of repeated sound segments in noise.

### Neural reconstruction of speech

To study whether speech is reliably represented in cortex in the presence of background noise, the temporal envelope of the underlying speech is reconstructed based on the neural responses to a speech-noise mixture (Fig. 2A). Neural reconstruction accuracy monotonically decreases with decreasing SNR for both native and foreign listeners. A 2-way repeated-measures ANOVA (listener group × SNR) is applied to the reconstruction accuracy, which shows significant main effects of the listener group (F_1,30_ = 4.22, P = 0.049) and SNR (F_3,90_ = 142.41, P < 0.001) and also the interaction between the 2 factors (F_3,90_ = 4.98, P = 0.003). At +9 dB and −6 dB SNRs, the reconstruction accuracy is higher for foreign than native listeners (+9 dB: P < 0.001, and −6 dB: P < 0.001, bootstrap, FDR corrected). If the relationship between reconstruction accuracy and SNR is fitted by a line. The slope of the line is significantly steeper for foreign listeners than native listeners (P < 0.001, bootstrap). Therefore, the neural reconstruction accuracy is more sensitive to noise in foreign listeners than native listeners.

**Fig. 2.**
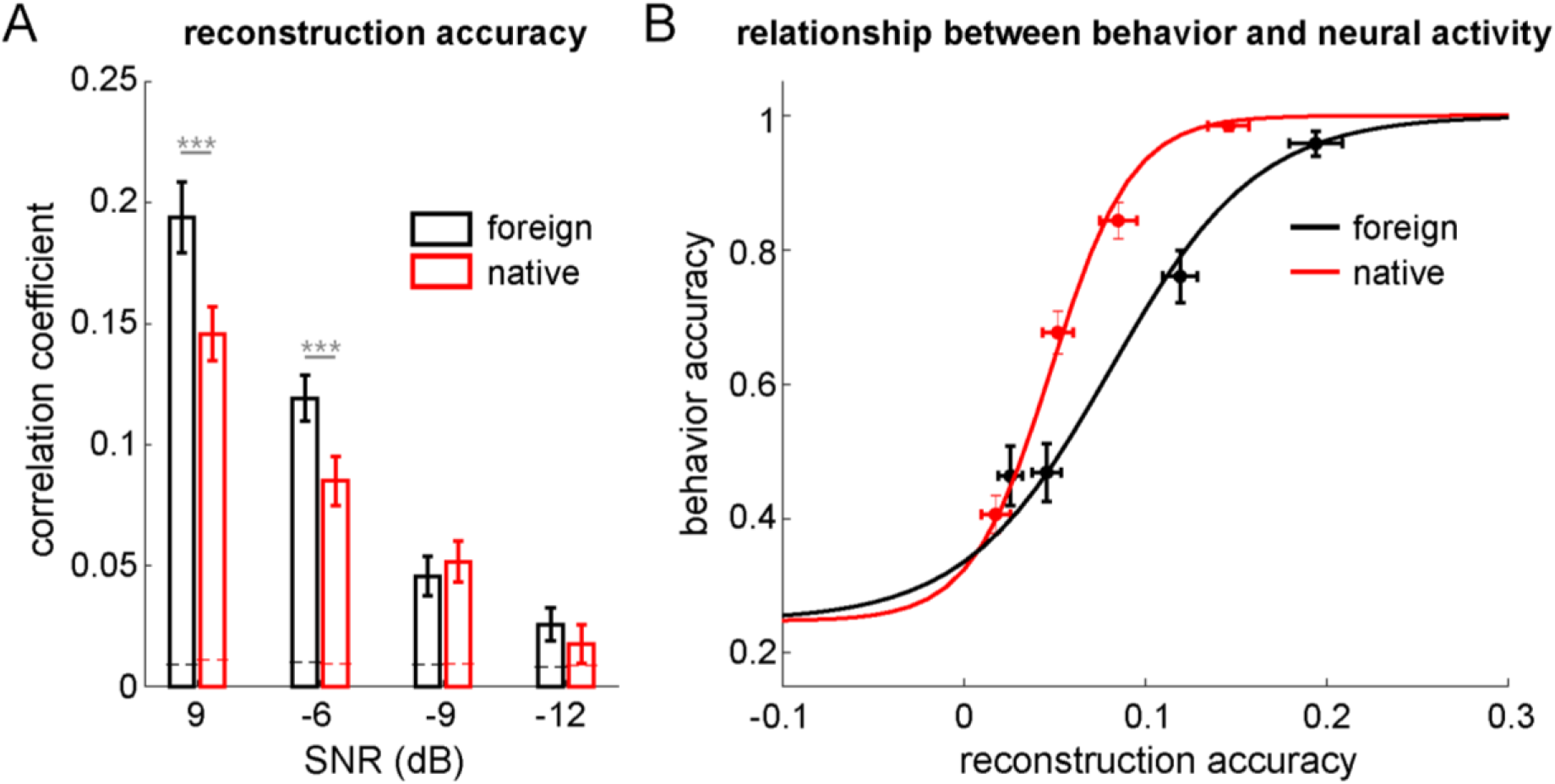
Neural reconstruction of speech envelope. (A) Reconstruction accuracy for native (red) and foreign listeners (black) under each SNR condition. Dashed lines denote the chance level. ***P < 0.001 (bootstrap, FDR corrected) (B) Relationship between behavioral accuracy and the neural reconstruction accuracy. Results from the 4 SNR conditions are shown and the error bars represent 1 SEM over listeners. The relationship between neural and behavioral accuracy is fitted by a sigmoid function for each group of listeners.

The relationship between behavioral accuracy and neural reconstruction accuracy is shown in Fig. 2B and is fitted by a sigmoid function. The growth rate of the fitted sigmoid function, which decides the slope of the sigmoid function, is significantly higher for native than foreign listeners (P = 0.0234, bootstrap, FDR corrected). Furthermore, the fitted location parameter of the sigmoid function is smaller for native than foreign listeners (P = 0.0234, bootstrap, FDR corrected). Those results indicate that the performance of detecting repeated sound segments is not entirely determined by envelope-tracking neural activity but additionally modulated by language processing.

### Temporal response function (TRF)

The TRF characterizes the time course of neural activity evoked by a unit power increase in the stimulus (Ding and Simon, 2012b). We first analyze the TRF derived from the actual stimulus, i.e., the speech-noise mixture, which reflects how the brain encodes the sound input. The predictive power averaged over listeners is shown in Fig. 3. The predictive power describes how accurately the speech envelope is tracked in each EEG channel. The topography of TRF predictive power shows a centro-frontal distribution when it can be reliably estimated, i.e., at +9, −6, and −9 dB SNRs. A 2-way repeated-measures ANOVA shows significant a main effect of SNR (F_3,90_ = 80, P < 0.001) and the interaction between SNR and listener groups (F_3,90_ = 5.85, P = 0.001). The main effect of listener groups, however, is not significant (F1,30 = 3.03, P = 0.092).

**Fig. 3.**
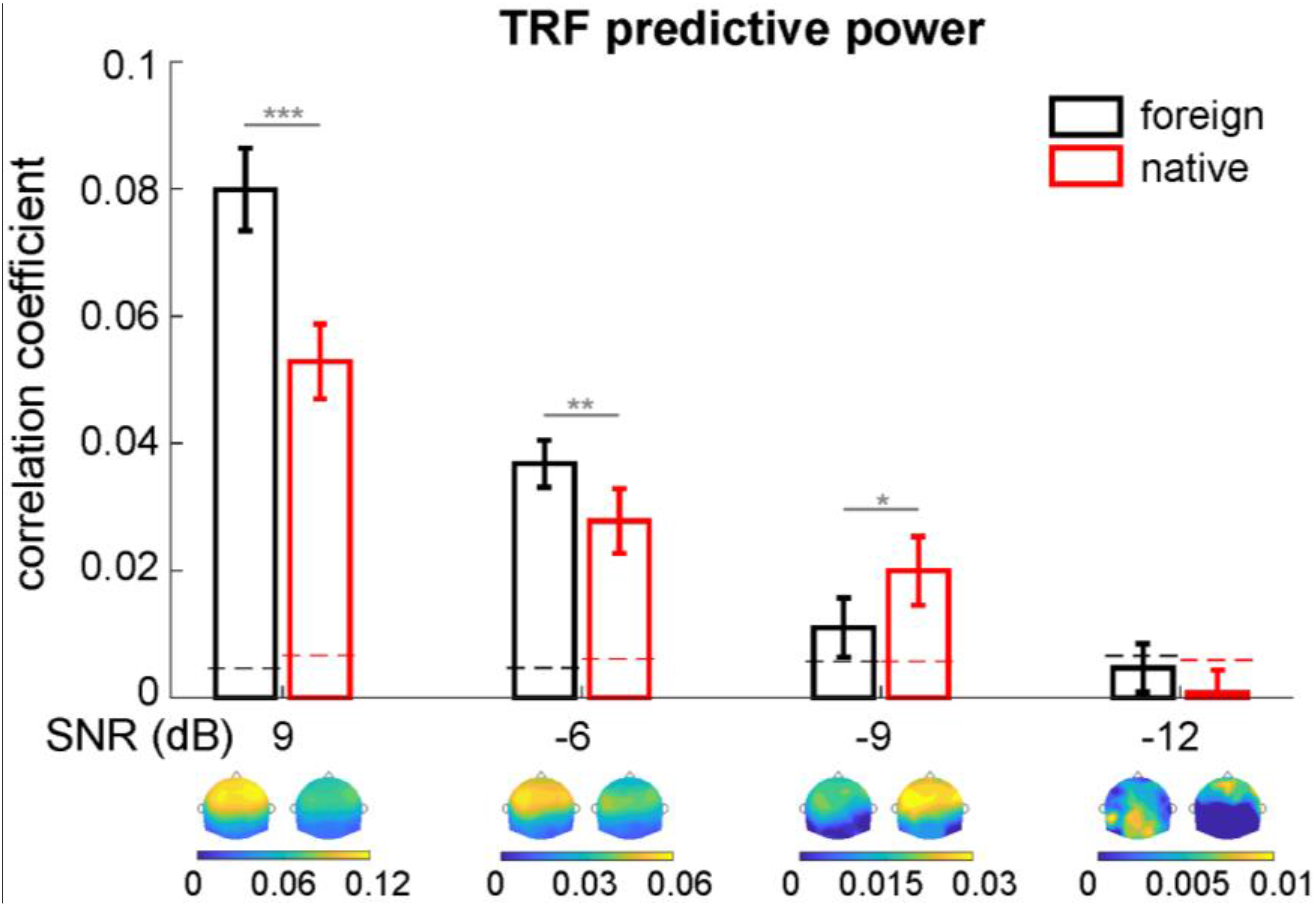
TRF predictive power for native (red) and foreign (black) listeners. The bar graph shows the predictive power averaged over all channels and the topography of the predictive power is shown underneath. Dashed lines indicate the 95% confidence interval of chance level. Error bars represent 1 SEM over listeners. *P < 0.05, **P < 0.01, ***P < 0.001, (bootstrap, FDR corrected).

Similar to neural reconstruction accuracy, the TRF predictive power is significantly higher for foreign listeners at +9 dB (P < 0.001, bootstrap, FDR corrected) and −6 dB (P = 0.003, bootstrap, FDR corrected) SNRs. At −9 dB SNR, however, the predictive power is higher for native listeners (P = 0.033, bootstrap, FDR corrected). If the relationship between predictive power and the SNR is fitted by a line, the slope of the line is shallower for native listeners (P < 0.001, bootstrap), suggesting that intelligibility enhances the robustness of envelope tracking in noisy environments.

To test if language knowledge and stimulus SNR modulate the spatial distribution of TRF predictive power, a classification analysis is employed to distinguish the 2 groups of listeners and 4 levels of SNRs based on the topography of predictive power. The predictive power from the 64 channels was first reduced to 2 dimensions using the LDA. The 2 groups of listeners are well separated at high SNRs in the first two LDA dimensions (Fig. 4A). The first 2 LDA dimensions approximate the 64-channel predictive power as the weighted sum of 2 topographic patterns, which are shown in Fig. 4B. When classifying the 64-channel predictive power into 8 categories (2 listener groups x 4 SNR levels), the classification accuracy is 38.3%, significantly higher than the chance-level performance of 12.5% (binomial test, P < 0.001). The accuracy to distinguish the 2 listener groups is 63.3% (pooling together SNR levels), higher than the chance performance of 50% (binomial test, P = 0.002), and the accuracy to distinguish the 4 SNR conditions is 56.3% (pooling together listener groups), also higher than the chance performance of 25% (binomial test, P < 0.001). The confusion matrix of the classification results is shown in Fig. 4C, which shows the decoded category of the data from each SNR condition for each group of listeners.

**Fig. 4.**
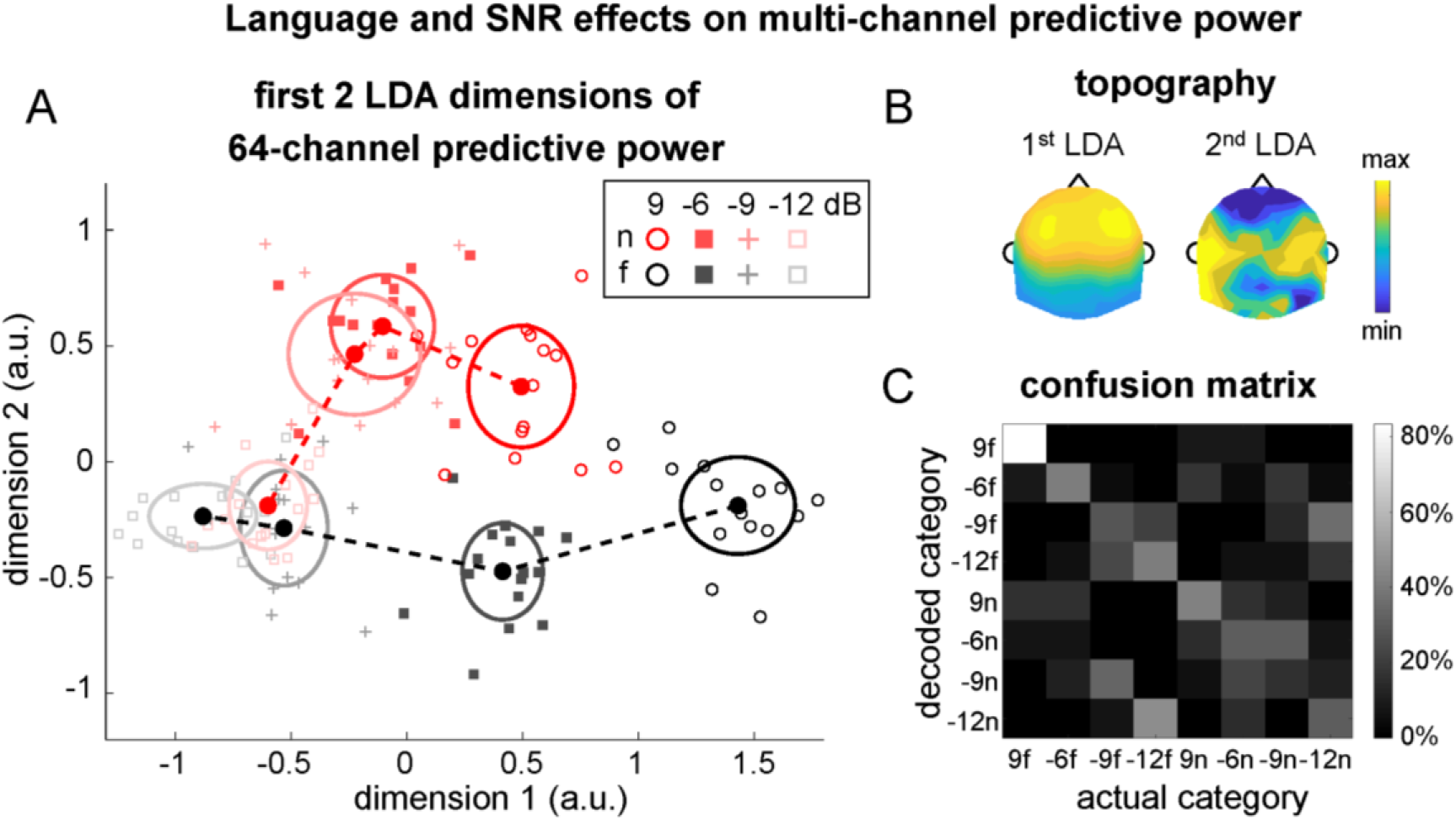
Classifying listener groups and SNR levels based on the topography of the predictive power. (A) Scatterplot of the first 2 LDA dimensions of the 64-channel predictive power. Lighter colors denote lower SNRs. Each marker represents data from 1 listener. The center of each ellipse is the mean across listeners while and the radius is 1 SD over listeners. (B) Topography of the first 2 LDA dimensions. (C) Confusion matrix of the classification accuracy shows a diagonal structure, indicating above-chance classification performance (f: foreign listener; n: native listener).

The amplitude and time course of the TRF are analyzed in the following. The RMS of the TRF across channels (normalized by subtracting the pre-stimulus value) is shown in Fig. 5AB. A 2-way repeated-measures ANOVA reveals a main effect of SNR on the TRF total power (F_3,90_ = 10.16, P < 0.001). Neither the main effect of listener groups (F_1,30_ = 0.71, P = 0.406) nor the interaction between listener groups and SNR (F3,90 = 2.14, P = 0.101) is significant. The TRF total power significantly increases from 9 dB to −6 dB (foreign: P < 0.001, and native: P = 0.01, bootstrap, FDR corrected). For foreign listeners the response gain decreases from −6 dB to −9 dB (P = 0.003, bootstrap, FDR corrected) and for native listeners the response gain decreases from −9 dB to −12 dB (P = 0.01, bootstrap, FDR corrected). These results demonstrate that the neural response gain exhibits a U-shaped relationship with the stimulus SNR. When the noise is relatively weak, the neural response gain increases to compensate the loss of stimulus contrast. When the noise is too strong, however, the neural response gain stops increasing.

**Fig. 5.**
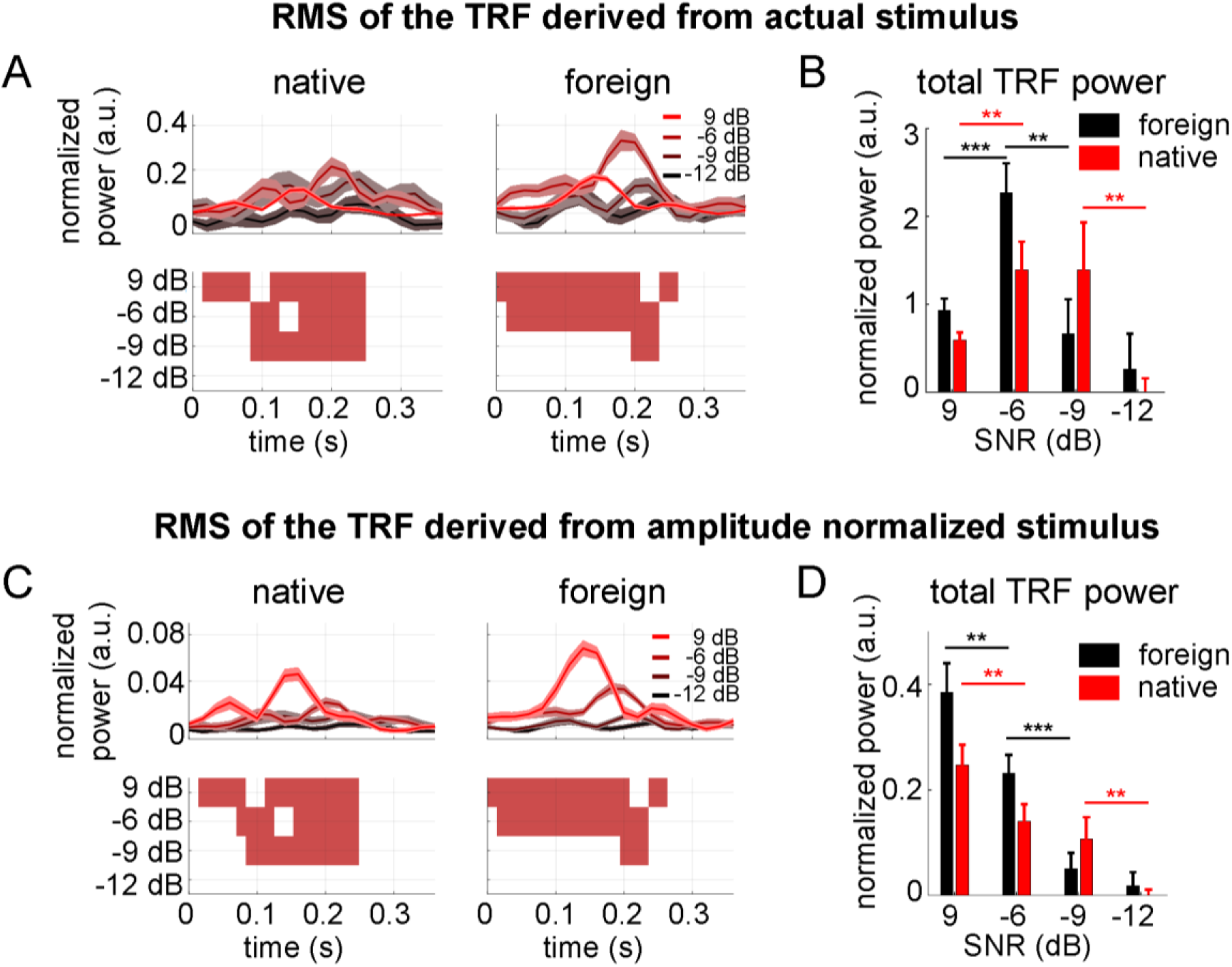
TRF derived from the actual stimulus (AB) or the amplitude-normalized stimulus (CD). (AC) The upper panel shows the TRF time course, i.e., the RMS over channels, and the shaded area denotes 1 SEM over listeners. The lower panel shows the time intervals in which the TRF amplitude is significantly higher than chance in red. Results from all SNR conditions are stacked vertically. (BD) Total power of the TRF. Error bar represents 1 SEM over listeners. *P < 0.05, **P < 0.01, (bootstrap, FDR corrected).

The previous analysis shows that the response gain depends on the stimulus SNR. In the following, we characterize how much the observed response gain change deviates from an ideal model in which the change in stimulus contrast is fully compensated. In this analysis, a new TRF is derived based on the amplitude-normalized stimulus envelope, i.e., the stimulus envelope divided by its standard deviation. If the change in neural response gain fully compensates the contrast reduction in the stimulus, the TRF derived from the amplitude-normalized stimulus will be independent of the stimulus SNR. The results, however, show that the TRF total power monotonically decreases with decreasing SNR, indicating that even between 9 dB and −6 dB, the change in response gain is not enough to fully compensate the loss of stimulus contrast.

In the following, a classification analysis is employed to quantify if language knowledge and the stimulus SNR modulate the spatial and temporal profiles of the TRF (Fig. 6). Using a procedure similar to that applied to classify the TRF predictive power, it is found that the listener groups and the SNR levels can be jointly classified with 45.3% accuracy (higher than chance, binomial test, P < 0.001). Since the predictive power analysis has already shown that language and SNR both modulate the spatial properties of the TRF, what remains unclear is whether the temporal information alone suffices to classify listener groups and stimulus SNRs. To address this issue, we remove the spatial dimension of the TRF by averaging the TRF over channels. Based on the channel-averaged TRF, classification is still above chance (43%, binomial test, P < 0.001), demonstrating that language knowledge and stimulus SNR modulate the TRF time course. In the classification analysis, the TRF model is derived from stimuli with normalized amplitude. For the TRF model derived from the actual stimulus, similar classification results are obtained based on the multi-channel TRF and the channel-averaged TRF (43% and 41.4% respectively, significantly above chance, binomial test, P < 0.001).

**Fig. 6.**
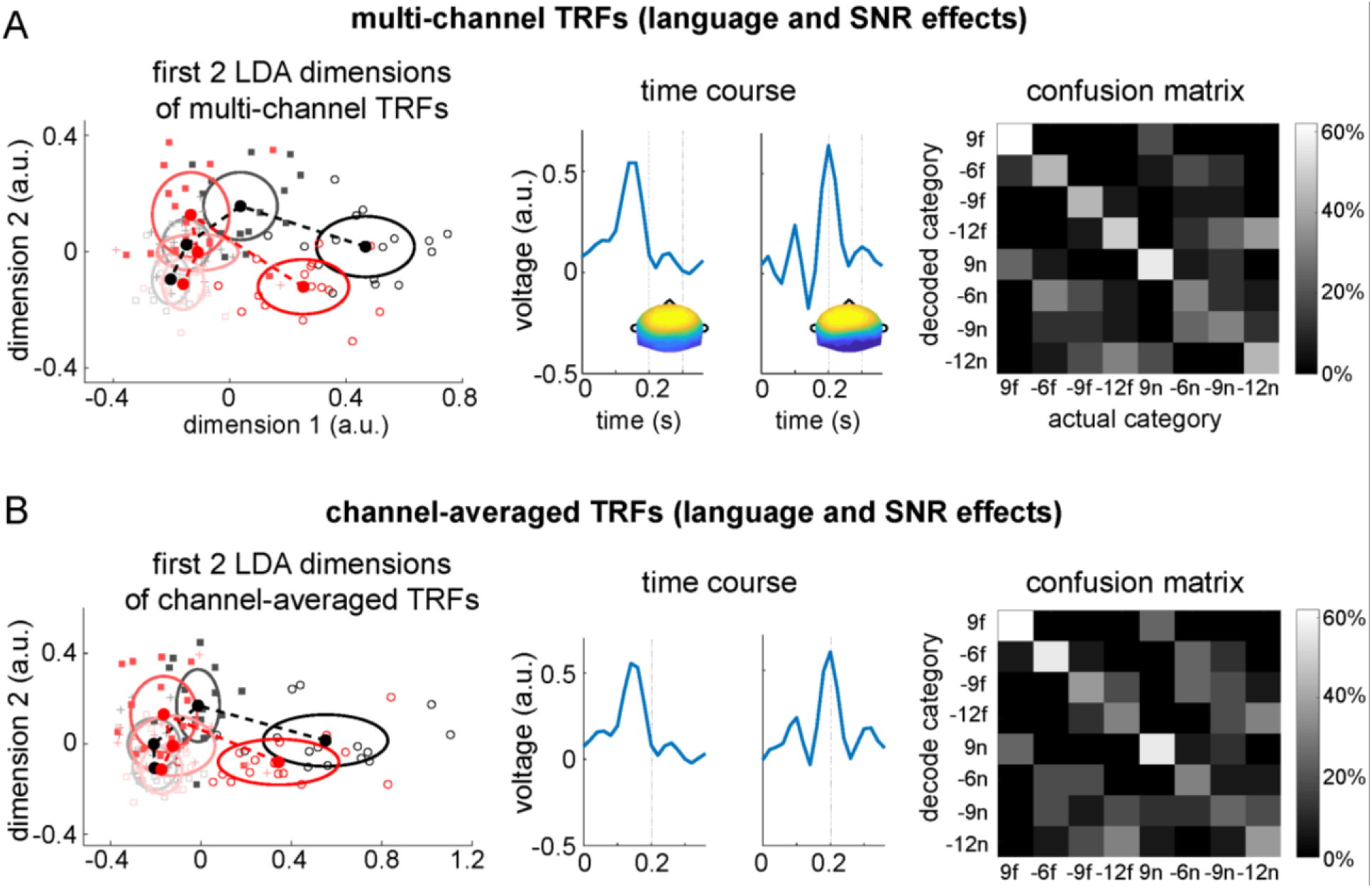
Classifying listener groups and SNR levels based on the multi-channel TRF (A) or channel-averaged TRF (B). The left panel shows the scatterplot of individual data for native (red) and unfamiliar (black) listeners. The central panels show the spatial and temporal profiles of the first 2 LDA dimensions. The right panels show the confusion matrix for the classification.

To further investigate whether the EEG response encodes the envelope of the actual stimulus, i.e., a speech-noise mixture, or the envelope of the underlying clear speech, we drive another TRF to describe the relationship between the EEG response and the underlying clear speech. As is shown in Fig. 7A, with decreasing SNR, the power of this TRF monotonically decreases. For the TRF total power, a 2-way repeated-measures ANOVA reveals a significant main effect of SNR (F3,90 = 34.85, P < 0.001) and a significant interaction between listener group and SNR (F3,90 = 2.99, P = 0.035), but no significant main effect of listener groups (F1,30 = 2.15, P = 0.153). Additionally, the normalized power is significant higher for foreign listeners at +9 dB (P = 0.034, bootstrap, FDR corrected) and −6 dB (P = 0.007, bootstrap, FDR corrected) when comparing with native listeners.

**Fig. 7.**
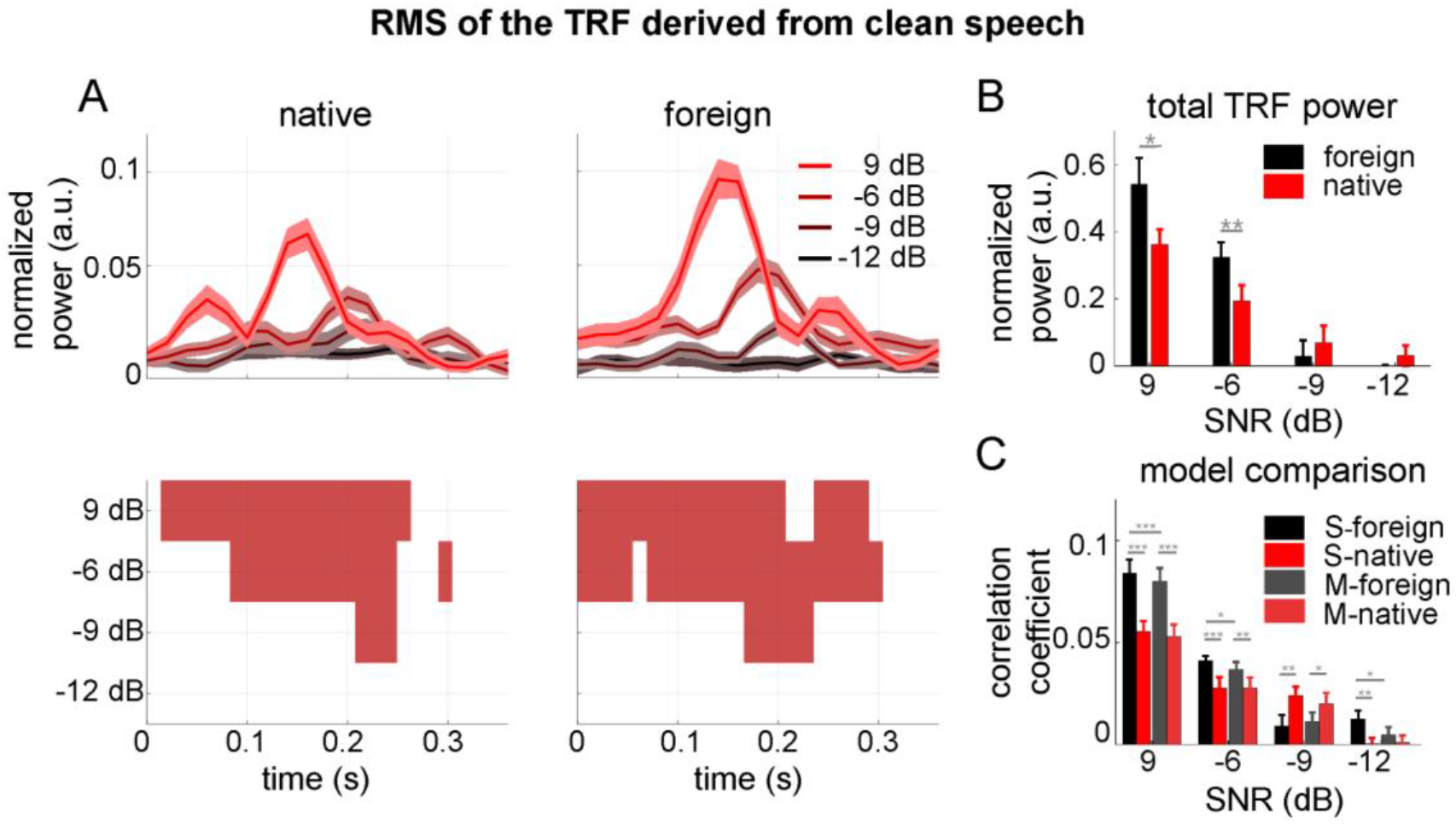
TRF derived from the underlying clear speech. (A) RMS of the TRF across channels (upper panels) and time intervals in which the TRF amplitude is significantly higher than chance (red areas, lower panels). The shaded areas in upper panels denote 1 SEM over listeners. (B) Normalized total power of the TRF. (C) Predictive power of the TRF derived from underlying clear speech (S) and the speech-noise mixture (M). Error bars represent 1 SEM over listeners. *P < 0.05, **P < 0.01, ***P < 0.001, (bootstrap, FDR corrected).

Finally, we analyze whether the actual stimulus or the underlying clean speech provides is better at modeling the EEG responses. Figure 7C compares the predictive power of the two models, which are comparable for native listeners. For foreign listeners, however, the model derived from clean speech yields slightly better predictive power at +9 dB (p < 0.001, bootstrap, FDR corrected) and −6 dB (p = 0.038, bootstrap, FDR corrected). This results indicates that even for an unknown language, the brain can actively reduce background noise and selectively encode the speech signal.

## Discussion

The temporal envelope is an acoustic feature that strongly contributes to speech intelligibility. This study demonstrates that the cortical representation of speech envelope is modulated by both auditory and language mechanisms, when the listeners perform a low-level auditory task that does not require speech comprehension. It is shown that envelope-tracking cortical activity can be generated based on domain-general auditory mechanisms and auditory gain control mechanisms can lead to a noise-robust representation even for unintelligible speech. Nevertheless, language processing does play a modulatory role and affect the spatial and the temporal profiles of speech-tracking activity.

### Auditory mechanisms underlying noise-robust speech representations

When speech is embedded in noise, the dynamic range of its intensity fluctuations, i.e., the intensity contrast, is compressed (Fig. 1B). If auditory cortex passively follows the stimulus envelope, the envelope-tracking response will reduce in amplitude when the noise level increases. Nevertheless, a number of studies have shown that a reduction in intensity contrast can be compensated at various auditory processing stages (Dean et al., 2005; Robinson and McAlpine, 2009), so that in auditory cortex neural activity is only weakly affected by the intensity contrast of the stimulus (Rabinowitz et al., 2011). MEG studies in humans show that neural tracking of the speech envelope is robust to noise and is barely influenced by the intensity contrast of the stimulus when the SNR is above ~-6 dB and the speech intelligibility is above ~50% (Ding and Simon, 2013). Robust neural encoding of the speech envelope has been attributed to auditory gain control mechanisms. Here, it is demonstrated that such mechanisms do not need feedback from high-level language processing and can be applied to unintelligible speech (Fig. 5A).

### Language processing contributes to robust speech processing in noise

High-context sentences can be better understood in noisy environments than low-context sentences or random words (Miller et al., 1951). Furthermore, prior knowledge about the content of a sentence can help the listeners to understand it in noise (Helfer and Freyman, 2005; Jones and Freyman, 2012; Kidd Jr et al., 2014; Yang et al., 2007). In the present study, we further demonstrate that language knowledge can even facilitate a low-level auditory task, i.e., detecting repeated sound patterns, at an intermediate SNR levels, i.e., −6 dB and −9 dB (Fig. 1C). The behavioral results are consistent with our neurophysiological findings that the precision of envelope-tracking activity, characterized by either the neural reconstruction accuracy (Fig. 2A) or the predictive power (Fig. 3), decreases faster for foreign listeners than native listeners. The current study shows that long-term language knowledge can influence cortical tracking of speech envelope and a recent study shows that recent exposure to a speech utterance can facilitate neural tracking of the envelope of that utterance in challenging listening environments (Wang et al., 2018). Taken together, the behavioral and neural results from the current study and other recent studies show that prior knowledge, either long-term language knowledge or recent listening experience, can modulate neural encoding of acoustic features, e.g., the speech envelope, and can boost the performance on low-level acoustic tasks that do not require speech comprehension.

### Influences of task and attention on neural tracking of speech envelope

Task and attention differentially influence envelope-tracking activity in different listening environments. When the listening environment contains multiple auditory streams, e.g., multiple speakers, selective attention strongly modulates cortical tracking of each auditory steam (Ding and Simon, 2012a; Kerlin et al., 2010; Mesgarani and Chang, 2012; O’Sullivan et al., 2014; Zion Golumbic et al., 2013). In a quiet listening environment, however, attention does not strongly affect neural tracking of the speech envelope. One study finds no overall change in speech tracking activity, characterized by the cross-correlation between EEG and speech envelope, when comparing an active listening condition with a condition in which the listener watch a silent movie (Kong et al., 2014). When comparing the same two conditions, another study in which the stimulus is an isochronous syllable sequence finds a subtle but statistically significant change in the power of envelope-tracking activity (Ding et al., 2018). In both these studies, the attention effects are much stronger when there are two competing speech streams. Task can influence attention allocation and can influence neural tracking of sound rhythms (Lakatos et al., 2013). However, during speech listening, when the task changes from a lexical task to a low-level speaker gender detection task, neural tracking of words is significantly reduced but neural tracking of the speech envelope is not significantly affected (Ding et al., 2018). In the current experiment, we employ a low-level auditory task that does not rely on speech comprehension. When the task changes, we expect the envelope-tracking response to be only weakly affected in relatively quiet listening environments. Future studies are needed, however, to investigate how envelope-tracking activity is influenced by task when the intensity of background noise is high.

### Intelligibility and cortical envelope tracking: stimulus manipulations

The relationship between envelope tracking activity and speech intelligibility has been extensively studied using 2 approaches. One approach varies speech intelligibility by manipulating acoustic features or the linguistic context, while the other approach studies individual differences in speech intelligibility. When the acoustic properties of speech is manipulated, inconsistent results have been reported in the literature about whether higher speech intelligibility is associated with more precise envelope tracking, especially in quiet listening environments. Some studies have reported no overall reduction in the precision of envelope-tracking activity when speech is played backward (Howard and Poeppel, 2010; Zoefel and VanRullen, 2016) or when the speech fine structure is corrupted (Ding et al., 2014). Nevertheless, other studies have found that the precision of envelope-tracking activity is reduced when speech is played backward (Gross et al., 2013; Park et al., 2015), when the spectro-temporal fine structure is corrupted (Luo and Poeppel, 2007; Peelle et al., 2013), or when the speech envelope is corrupted (Doelling et al., 2014).

In noisy listening environments, it has been more consistently reported that when an acoustic interference signal reduces speech intelligibility, it also reduces the precision of cortical envelope tracking (Ding and Simon, 2013; Kong et al., 2015; Vanthornhout et al., 2018). The current results also show decreased envelope tracking accuracy when the noise level increases. A potential issue for the acoustic manipulation approach, however, is that although it can demonstrate the correlation between speech intelligibility and envelope tracking, it cannot establish a causal relationship between them (Ding and Simon, 2014). A related method to manipulate speech intelligibility is to use the priming paradigm. In some conditions, an unintelligible sentence becomes intelligible if the listeners hear an intelligible version of the same sentence in advance. Using the priming approach, however, two studies have concluded that intelligibility does not modulate envelope tracking (Baltzell et al., 2017; Millman et al., 2015). Nevertheless, the priming paradigm cannot distinguish sensory-level adaptation effect caused by priming and the intelligibility changed caused by priming.

When the linguistic content of speech is manipulated, previous studies have shown comparable or reduced envelope-tracking for sentences than random syllables or words. When the linguistic structure is manipulated without changing the speech envelope, one study shows neural tracking of the speech envelope remains unchanged (Ding et al., 2016). In another study, when listening to a sequence of syllables are grouped into artificial words, neural tracking of the syllabic-rate speech envelope is reduced after the listeners learn the words via statistical learning (Buiatti et al., 2009). The previous two studies discussed here both presented speech syllables isochronously.

Using natural utterances, one study observes more accurate delta-band envelope tracking for sentences constructed by pseudowords, compared with sentences constructed by real words (Mai et al., 2016). One explanation for this phenomenon is that linguistic context provides additional cues for speech comprehension so that the brain relies less on acoustic cues in the speech envelope. Another explanation is that neural tracking of the linguistic content (Ding et al., 2016) competes with neural tracking of envelope and therefore reduces envelope-tracking activity. These two explanations are also potential reasons why better behavioral performance is achieved with lower envelope tracking precision for native listeners (Figs. 2B & 3).

### Intelligibility and cortical envelope tracking: individual differences

The relationship between envelope tracking activity and intelligibility can also be characterized by individual differences. One study has compared the neural responses of native Italian and Spanish listeners while they listen to Italian, Japanese, and Spanish utterances (Peña and Melloni, 2012). It is shown that speech intelligibility does not change in low-frequency power, a gross measure that can reflect both envelope-tracking activity and other neural responses. The current study extends this previous study by explicitly analyzing neural tracking of the speech envelope. In another study, when listening to sentences constructed by isochronously presented syllables, the strength of the envelope-tracking response is comparable for native and foreign listeners (Ding et al., 2016). The current study, however, find that neural tracking of the speech envelope is more precise for non-native listeners for utterances with a natural rhythm.

Within the population of native listeners, individuals vary in their ability to understand speech in noisy environments. In challenging listening conditions, studies have consistently shown that individuals showing more precise envelope-tracking activity tend to understand speech better (Ding et al., 2014; Ding and Simon, 2013; Doelling et al., 2014; Kong et al., 2015; Vanthornhout et al., 2018). In contrast, a more precise envelope-tracking response is observed for older listeners compared with younger listeners, in both quiet and noisy listening environments (Presacco et al., 2016), even though older listeners have more trouble understanding speech in noise. Across studies, it is suggested that speech-tracking activity is generally related to intelligibility at the individual level with in a relatively homogeneous population, e.g., young native listeners. It remains unclear, however, whether this effect is driven by individual differences in auditory ability or language ability. Across populations, the relationship between intelligibility and envelope tracking precision seems to be heterogeneous.

Taken together, the current results and previous studies (Buiatti et al., 2009; Mai et al., 2016; Presacco et al., 2016) demonstrate that when the stimulus acoustic properties are controlled, language processing generally reduces the precision of envelope-tracking activity in quiet listening environments. Critically, however, we find that the precision of envelope tracking activity is differentially modulated by speech intelligibility in different EEG channels so that the topography of the TRF predictive power can be used to distinguish the native and foreign listeners (Fig. 4). Similarly, the time course of the TRF, i.e., the RMS over channels, is also sufficient to distinguish the two groups of listeners (Fig. 6). These results suggest that future research needs to consider the spatio-temporal dynamics of neural activity when discussing how intelligibility modulates envelope-tracking responses.

## Acknowledgement

Work supported by National Natural Science Foundation of China 31500873 (ND), 31771248 (ND), Zhejiang Provincial Natural Science Foundation of China LR16C090002 (ND), and research funding from the State Key Laboratory of Industrial Control Technology, Zhejiang University (ND).

